# UNC-45A is Highly Expressed in the Proliferative Cells of the Mouse Genital Tract and in the Microtubule-Rich Areas of the Mouse Nervous System

**DOI:** 10.1101/2021.03.19.436218

**Authors:** Valentino Clemente, Asumi Hoshino, Joyce Meints, Mihir Shetty, Tim Starr, Michael Lee, Martina Bazzaro

**Author notes:** Corresponding author: Martina Bazzaro, Masonic Cancer Center, Room 490, 420 Delaware Street S.E, Minneapolis, Minnesota 55455. Equal contributors (Valentino Clemente, Asumi Hoshino).

## Abstract

UNC-45A is a cytoskeletal-associated protein with a dual and non-mutually exclusive role as a regulator of the acto-myosin system and as a Microtubule (MT)-destabilizing protein. UNC-45A is overexpressed in human cancers including in ovarian cancer patients resistant to the MT-stabilizing drug Paclitaxel. Mapping of UNC-45A in the mouse upper genital tract and central nervous system reveals its enrichment in highly proliferating and prone to remodeling cells and in microtubule-rich areas of in the ovaries and in neurons respectively. In both apparatuses UNC-45A is also abundantly expressed in the ciliated epithelium. Because regulators of acto-myosin contractility and MT stability are essential for the physiopathology of the female reproductive tract and of neuronal development our findings suggest that UNC-45A may have a role in ovarian cancer initiation and development and in neurodegeneration.

## Introduction

UNC-45A is a member of the UCS (UNC-45/CRO1/She4p) protein family (*1, 2*) with a dual and non-mutually exclusive role as a regulator of the acto-myosin system (*3, 4*) and as a MT-destabilizing protein (*5–7*). As a key regulator of cytoskeletal activities UNC-45A participates in a number of cellular functions including cytokinesis (*8–10*), exocytosis (*11*), and axonal growth (*12*). UNC-45A is overexpressed in breast and ovarian cancer as compared to their normal counterpart (*8–10*) and in ovarian cancer patients that are resistant to the microtubule (MT)-stabilizing drug paclitaxel in ovarian cancer (*6*). Regulators of acto-myosin contractility and MT stability are essential for both ovarian cancer cells proliferation and neuronal development. For instance, dysregulation of the Rho/ROCK signaling pathway is commonly found in ovarian cancer (*13–16*) and implicated in the pathophysiology of nervous system (*17, 18*) (*19, 20*) (*21*) (*22*). A number of MT-destabilizing proteins are also expressed in both neurons (*23–27*) and cancer cells including ovarian cancer cells (*28–30*) where they play roles spanning from regulating symmetrical and asymmetrical cell division (*31*) to regulate MT mass (*6*) to regulate sensitivity to MT-targeting agents (*6, 32*).

In this study we investigated the UNC-45A expression pattern in the mouse upper genital tract and in the brain. In the ovaries and fallopian tube, we found that UNC-45A is enriched in highly proliferating and prone to remodeling cells. In the brain we found that UNC-45A is expressed in the microtubule-rich regions of the mouse central nervous system and in the mouse nerve roots. We also found that UNC-45A is abundantly expressed in the cilia of cells in both the upper genital tract and the brain. Taken together these findings suggest that UNC-45A may play a role in the physiology and pathology of the female reproductive apparatus and of the central nervous system.

## Results

### UNC-45A is expressed in the mouse upper genital track with a stronger expression in proliferating cells and in cilia

We and others have previously shown that UNC-45A is expressed in human ovarian surface epithelial cells and in human breast cells and overexpressed in highly proliferative cells of ovarian and breast cancers as compared to their stoma counterpart (*8, 9*). Here, we performed H&E and UNC-45A staining (Figure 1A, *upper and lower panels* respectively) in mouse ovaries and fallopian tube to determine UNC-45A expression pattern in the mouse upper genital tract. We found that UNC-45A is abundantly expressed in cells of the early follicles (Figure 1B, **i**), the corpus luteus (Figure 1B, **ii**) and the later follicles (Figure 1B, **iii**). Furthermore, UNC-45A is strongly expressed in oocytes (Figure 1B asterisk) and in the mouse surface epithelium (Figure 1B, *inset, arrow*) while the stroma is mostly negative. In the fallopian tubes, UNC-45A is weakly expressed in the smooth muscle cells and more intensively expressed in the fallopian tube epithelium an in the particular in the cilia of the fallopian tube epithelium (Figure 1C and its inset *green arrows*). Importantly cilia are microtubule (MT)-based structures capable of moving themselves using dynein contractility and we have recently shown that UNC-45A is a Microtubule-Associated-Protein (MAP) responsible for modulating MT dynamics (*33*) (*7*) (*34*)). Taken together this suggests that in the mouse upper genital tract UNC-45A expression is stronger in highly proliferative and prone to remodeling cells and that in that UNC-45A may play a role in regulating cilia functions.

**Figure 1.**
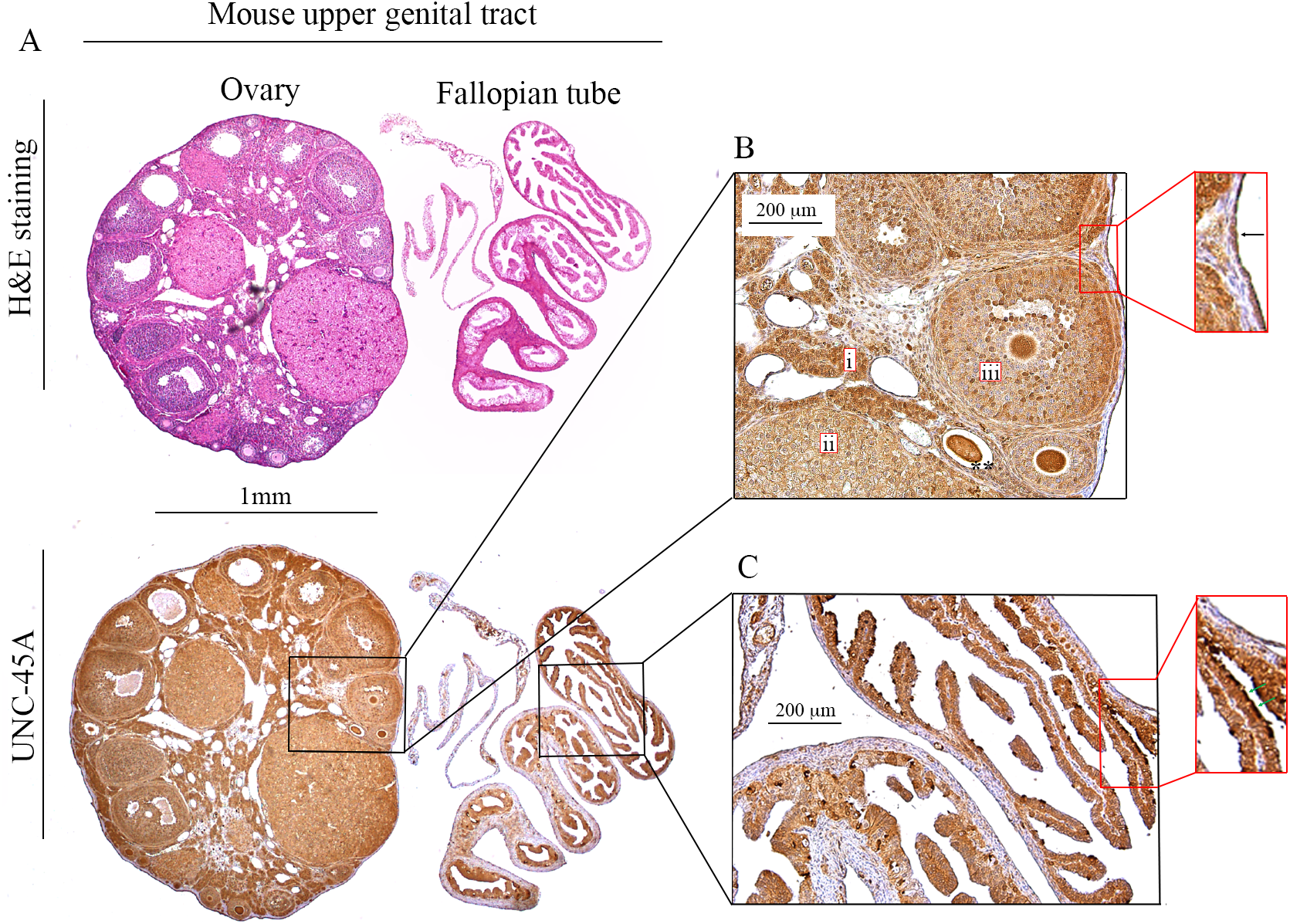
UNC45A expression in mouse ovaries and fallopian tube. **A.** H&E (*upper panel*) and UNC-45A (*lower panel*) staining of the mouse ovary and fallopian tube. **B.** Details of mouse ovary with follicular structures strongly stained for UNC-45A (**i**. early follicles, **ii**. corpus luteus, and **iii**. later follicles) and containing oocytes (asterisks). *Inset*, mouse ovarian surface epithelium (*arrow*) positive for UNC-45A while the stroma is negative. **C.** Details of mouse fallopian tube with epithelium strongly stained for UNC-45A. *Inset*, cilia in fallopian tube epithelium (green arrows) strongly positive for UNC-45A.

### UNC-45A is expressed in the microtubule-rich regions of the mouse central nervous system (CNS) and in the mouse nerve roots

We have recently shown that UNC-45A is a Microtubule-Associated-Protein (MAP) with MT-severing properties (***5–7***), which is important for the neurite development (***12***). In neurons, MTs play a pivotal role for both the structure and function of axons and dendrites, whereas MAPs have been extensively shown to be major regulators of the physiology and pathology of the nervous system (***35, 36***). Here, we performed H&E and UNC-45A staining (Figure 2A and Figure 2B respectively) of whole adult mouse brain. H&E staining allowed for visualization of brain anatomical structures including the cerebellum (Figure 2A, **i**), the hippocampus (Figure 2A, **ii** and its inset) and the striatum (Figure 2A, **iii**). We found that in the cerebellum UNC45A is most strongly expressed in the white matter (Figure 2B, **i**, and its inset, ***red arrow***), while the intensity of the staining decreases progressively in the granular (Figure 2B, **i**, and its inset, ***green arrow***) and in the molecular (Figure 2B, **i**, and its inset, ***red asterisks***) layers of the cortex. This difference in UNC-45A staining intensity between the two cortical structures might be explained by their peculiar architecture: the granular layer is formed by neuronal bodies intercalated to the axonal fibers descending into the white matter, while the molecular layer by a sparse network of dendrites, some of which stand out from the background (Figure 2B, **i**, next to the asterisks) due to their high UNC45A expression. We also found that UNC-45A is abundantly expressed in the dendrites of the CA1 hippocampal neurons (Figure 2B, **ii**. and its inset, ***black arrow).*** In the striatum, UNC-45A is strongly expressed in the nervous bundles formed by axons crossing this grey structure (Figure 2B, **iii**, *black arrows)*. To gain more details on the pattern of UNC-45A expression in the mouse brain, we performed immunofluorescence analysis on the whole mouse brain stained for UNC-45A (green), the neuronal marker NeuN (***37***) (magenta) and DAPI (blue) to stain nuclei (Figure 3A). As shown in Figure 3B and C, in hippocampal CA1 neurons, UNC-45A has a perinuclear and cytoplasmic localization and is particularly enriched in the dendrites (yellow arrows). Noteworthy, UNC-45A staining is absent in oligodendrocytes, which can be recognized by the fact that they are DAPI positive, NeuN negative and surrounded by a black area represented by the myeline sheath (Figure 3C, ***yellow asterisk***). Furthermore, UNC-45A was also confirmed to have a strong staining in the nervous bundles of the striatum (Figure 3D, ***white asterisks***).

**Figure 2.**
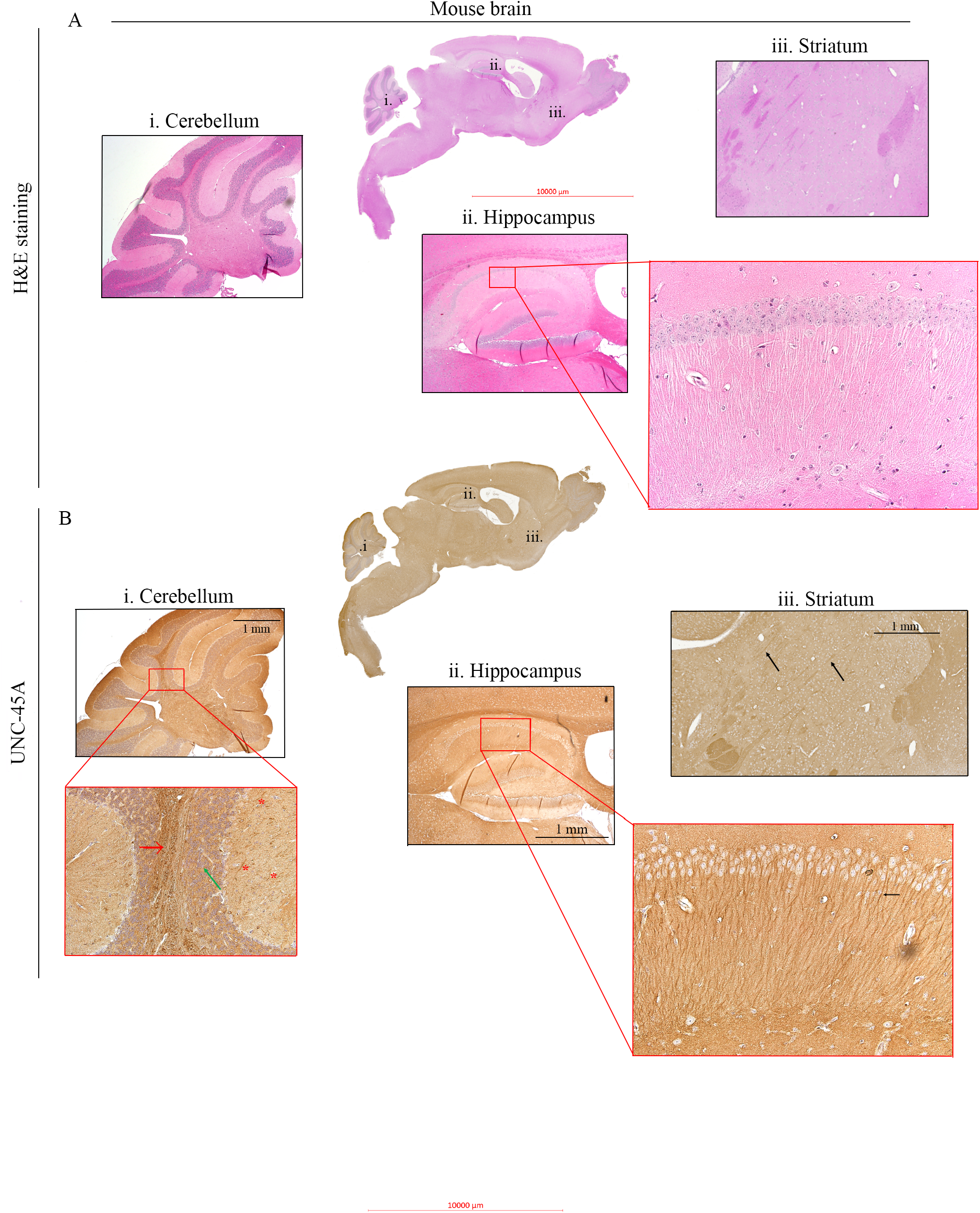
UNC45A expression in mouse brain via IHC. **A.** H&E staining of whole mouse brain. **i**. Cerebellum, **ii**. Hippocampus, **iii**. Striatum. **B.** UNC-54A staining in whole mouse brain. **i.** details on UNC-45A staining in Cerebellum (*inset*) from the center to the sides: white matter (red arrow) granular layer (*green arrow*), molecular layer (*red asterisks*), **ii.** details of UNC-45A staining in hippocampus, *inset* closer view on hippocampal CA1 neurons, with clearly visible dendrites (*black arrow*). **iii.** details of UNC-45A staining in striatum, arrows indicate stronger UNC-45A staining in nervous bundles.

**Figure 3.**
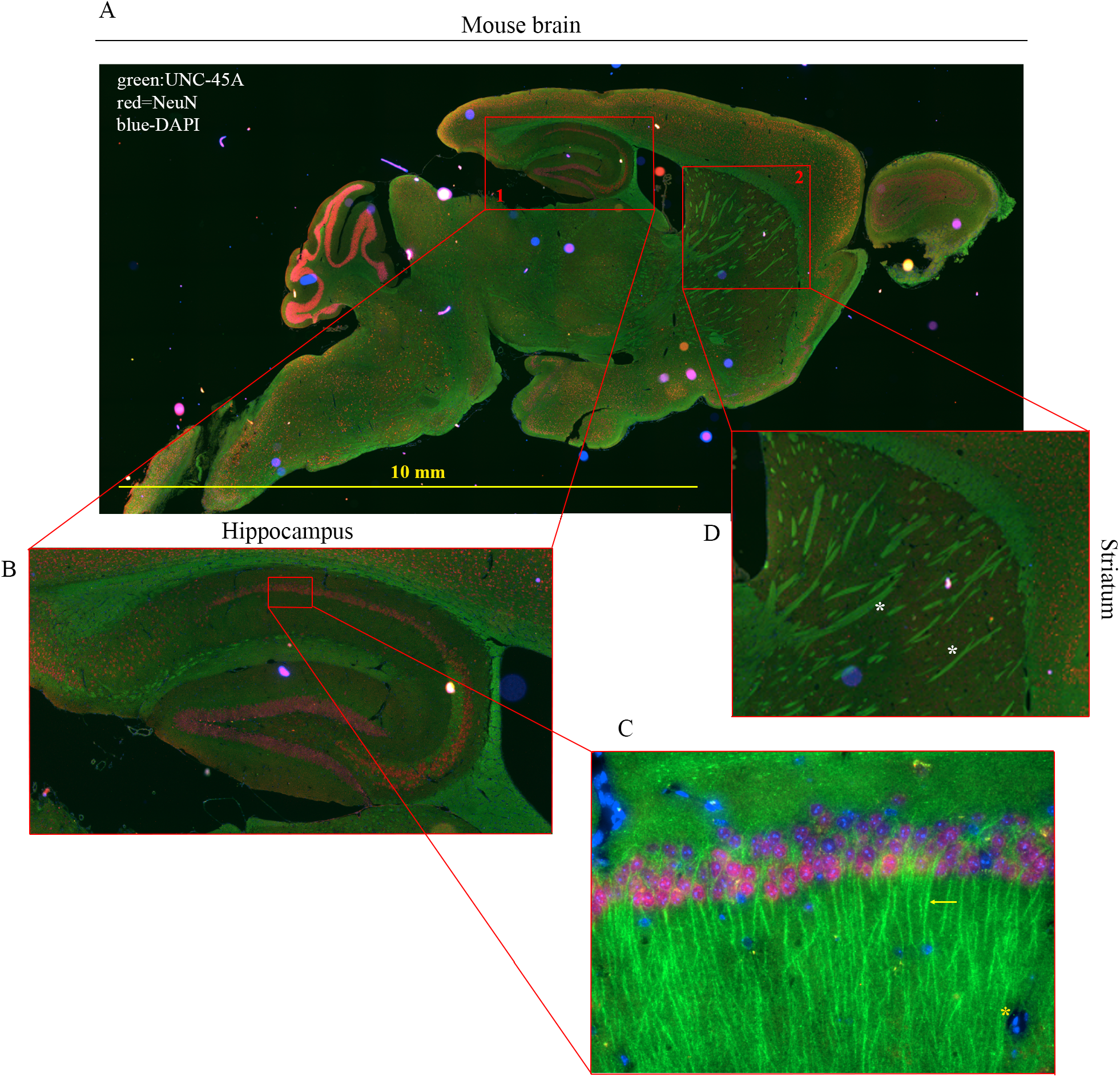
UNC45A expression in mouse brain via IF. **A.** Immunofluorescence staining for UNC-45A (green), NeuN (magenta) and DAPI (blue) in whole mouse brain. **B.** Details of stained hippocampus. **C.** Close-up of UNC-45A staining in axons of hippocampal neurons (yellow arrow), DAPI staining of nuclei and NeuN staining the neuronal soma. Details of absent UNC-45A staining in oligodendrocytes (*yellow asterisk*). **D.** Details of UNC-45A staining in the striatum (*white arrows* indicate the nervous bundles).

Next, we performed H&E and UNC-45A staining of the mouse spinal cord (Figure 4A upper and lower panels respectively) and mouse nerve root (Figure 4E, upper and lower panel respectively). In the spinal cord, UNC45A is more abundantly expressed in the white matter (Figure 4B, **i**.) than in the grey matter (Figure 4B, **ii**). In the white matter, intense UNC-45A staining can be seen in cross sections of single axons (Figure 4C), surrounded by UNC-45A-negative myeline sheath. Interestingly, a strong UNC-45A expression can also be found in the feet of the astrocytes forming the hematoencefalic barrier in the area around vessels (Figure 4B, *green arrow*), while the ependymocites express UNC45A prevalently in their cilia (Figure 4D, *red asterisk*), with the cytoplasm being just weakly positive. In the nerve root, the meninges (Figure 4E lower panel, *black arrow*) are negative for UNC-45A while a strong UNC-45A staining can be found in both, longitudinal (Figure 4F) and cross (Figure 4G) sections. In the cross section of the nerve roots, it is evident how strongly UNC-45A positive axons are surrounded by negatively stained myelin sheath (*red arrow*).

**Figure 4.**
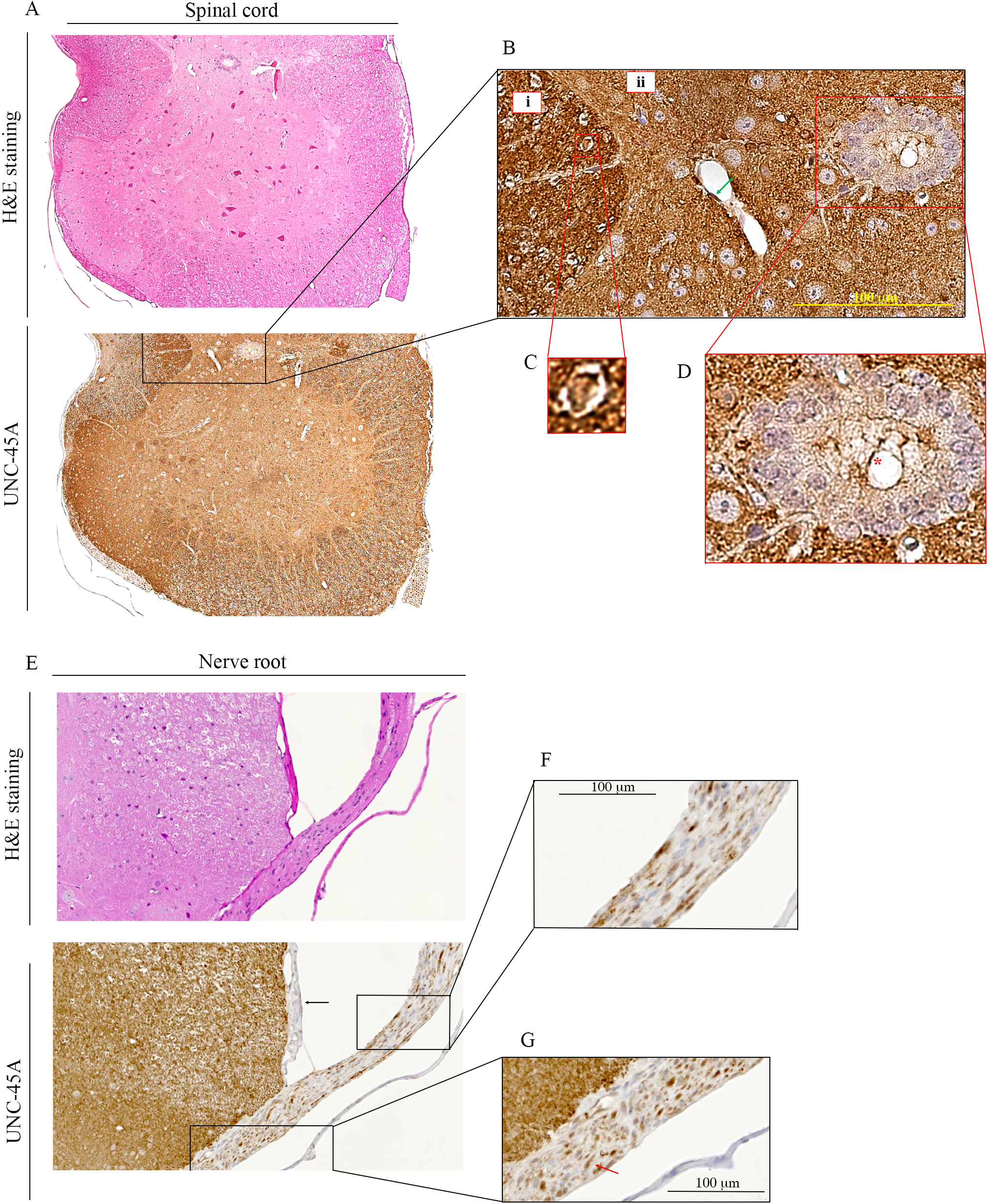
UNC-45A expression in mouse spinal cord and nerve root. **A.** H&E (*upper panel*) and UNC-45A (*lower panel*) staining of the mouse spinal cord. **B.** Details of UNC-45A staining in white matter (i), grey matter (ii), perivascular area (*green arrow*). **C.** Detail of UNC-45A staining of a single cross-sectioned axon with its myelin coat. **D.** UNC-45A staining in ependymocites and their ciliary structures (*red asterisk*). **E.** H&E (*upper panel*) and UNC-45A (*lower panel*) staining of the mouse nerve root (*black arrow* indicates the meninge). **F.** UNC-45A staining in longitudinal section of the nerve root. **G.** UNC-45A staining in cross section of the nerve root (*red arrow* indicates the myelin sheath).

Taken together, this suggests that UNC45A may play a relevant role in nervous system physiology. In particular, considering the crucial roles of microtubules and their MAPs in neuronal biology (*35, 36*), the preferential and abundant expression of UNC45A in axons and dendrites of mature neurons, shows that this protein is not only required for their development (*12*), but may also be important for the neuronal functions and homeostasis. This, could thus hint at a role for UNC45A in more complex processes such as brain plasticity and/or neurodegeneration.

## Discussion

Here we found that in the mouse upper genital tract UNC-45A is enriched in proliferating and prone to remodeling cells as compared to quiescent cells. These findings, along with the fact that UNC-45A is overexpressed in ovarian cancer (*9*) and that MT-severing proteins are involved in oocyte development (*38–40*), hint to the potential relevance of UNC-45A in pathophysiology of the ovaries including fertility and ovarian cancer initiation. We also found that UNC-45A is expressed in neurons in the mouse nervous system and is particularly enriched in areas containing axons and dendrites. Interestingly, Tau, a MAP which stabilizes MT via preventing MT severing (*41*) is largely expressed in brain, its hyperphosphorylation is associated with Alzheimer’s disease (AD) and other tauophathies (*42–47*), and is abnormally expressed in ovarian cancer (*48, 49*). Importantly, drugs targeting MT stability is an established anti-cancer approach for ovarian cancer (*50, 51*) and stabilization of neuronal MTs can attenuate neurodegenerative in mouse models of AD and other tauopathies (*52*). In this context, our results suggests that UNC-45A may play a role in the physiology of the central nervous system as well as in neurodegeneration. Lastly, we found that UNC-45A is abundantly expressed in the ciliated columnar epithelium of the fallopian tube and in the cilia of the ependymocytes. Interestingly, loss-of-function mutations in UNC-45A causes a syndrome characterized by diarrhea, cholestasis, bone fragility, and impaired hearing (*53*), all of which symptoms involve organs where cilia play important roles in physiology (*54–56*). In this context, our results suggests that UNC-45A may play a role in the physiology of ciliated epithelia as well as in ciliopathies.

## Materials and methods

### Sample preparation of mouse upper genital track

Ovaries and fallopian tube from adult C57BL/6 were preserved in 10% formalin buffer solution for 24 hours followed by 70% ethanol to prevent the tissue from decomposing and embedded in paraffin.

### Sample preparation of mouse brain and spinal cord

Brain and spinal cord from adult C57BL/6 were preserved in 10% formalin buffer solution for 24 hours followed by 70% ethanol to prevent the tissue from decomposing and embedded in paraffin.

### Antibodies

Anti-UNC-45A Rabbit Polyclonal antibody (19564-1-AP, Protein Tech) was used for IHC and IF. The specificity of this antibody has been previously validated by us and others by Western Blot, IF and IHC analyses (*5, 6*).The Anti-NeuN mouse monoclonal antibody (ThermoFisher MA5-33103) was used for IF. Secondary antibodies for IF analyses were FITC-conjugated Goat Anti-Rabbit IgG used at 1:200 dilution (Jackson ImmunoResearch Laboratories) and Texas-Red conjugated Goat Anti-Mouse IgG used at 1:200 dilution (Jackson ImmunoResearch Laboratories).

### Immunohistochemical staining of mouse upper genital track, brain and spinal cord

5 *μ*m formalin fixed; paraffin embedded (FFPE) tissue sections were subjected to hematoxylin & eosin staining and immunohistochemistry (IHC) for UNC-45A. Slides were deparaffinized with 100% xylene and rehydrated with gradient ethanol (100%, 95%, and 80%). Antigen retrieval was carried out with 1X Reveal Decloaker (Biocare medical) in a vegetable steamer for 30 min at 100°C. The slides were blocked by Background Sniper (BS966H, Biocare medical) for 13 min at room temperature. Sections were incubated with anti-UNC45A antibody at a dilution of 1:100 for mouse brain and spinal cord, 1:200 for mouse upper genital tract overnight at 4°C, followed by Biotin-SP-conjugated AffiniPure Goat Anti-Rabbit IgG at a dilution of 1:200 for 30 min at room temperature and incubation with horseradish peroxidase streptavidin at a dilution of 1:125 (405210, BioLegend) for 30 min at room temperature. After the staining was developed with 3,3’-diaminobenzidine (926506, BioLegend) for 3 min, the slides were counterstained with Harris hematoxylin.

### Brighfield imaging

Brightfield imaging were acquired with a Zeiss Axio Scan.Z1 system (Zeiss, German) at 40X magnification. Images of details were taken with a BX40 light microscope, DP72 camera, and cellSens Standard v1.16 imaging software (Olympus, Japan).

### Immunofluorescence Microscopy

For immunofluorescence analysis of UNC-45A and NeuN in mouse brain, 5 *μ*m formalin fixed; paraffin embedded (FFPE) tissue sections were deparaffinized, rehydrated, retrieved and blocked similarly to what described in the immunohistochemical staining section. Sections were then incubated with anti-UNC45A and anti-NeuN at the indicated concentration overnight and 4 degrees. Secondary antibodies were used at the indicted concentration for 1 hour at room temperature and sections were mounted with mounting media containing DAPI (F6057 Sigma-Aldrich). Images were obtained on an Axiovert 200 microscope (Zeiss, Thornwood, NY) equipped with a high-resolution CCD camera.

